# Machine learning reveals heterogeneous responses to FAK, Rac, Rho, and Cdc42 inhibition on vascular smooth muscle cell spheroid formation and morphology

**DOI:** 10.1101/2020.01.30.927616

**Authors:** Kalyanaraman Vaidyanathan, Chuangqi Wang, Amanda Krajnik, Yudong Yu, Moses Choi, Bolun Lin, Su-Jin Heo, John Kolega, Kwonmoo Lee, Yongho Bae

**Author notes:** These authors contributed equally: Kalyanaraman Vaidyanathan and Chuangqi Wang.

## Abstract

Atherosclerosis and vascular injury are characterized by neointima formation caused by the aberrant accumulation and proliferation of vascular smooth muscle cells (VSMCs) within the vessel wall. Understanding how to control VSMCs would advance the effort to treat vascular disease. However, the response to treatments aimed at VSMCs is often different among patients with the same disease condition, suggesting patient-specific heterogeneity in VSMCs. Here, we present an experimental and computational method called HETEROID (Heterogeneous Spheroid), which examines the heterogeneity of the responses to drug treatments at the single-spheroid level by combining a VSMC spheroid model and machine learning (ML) analysis. First, we established a VSMC spheroid model that mimics neointima formation induced by atherosclerosis and vascular injury. We found that FAK-Rac/Rho, but not Cdc42, pathways regulate the VSMC spheroid formation through N-cadherin. Then, to identify the morphological subpopulations of drug-perturbed spheroids, we used an ML framework that combines deep learning-based spheroid segmentation and morphological clustering analysis. Our ML approach reveals that FAK, Rac, Rho, and Cdc42 inhibitors differentially affect the spheroid morphology, suggesting there exist multiple distinct pathways governing VSMC spheroid formation. Overall, our HETEROID pipeline enables detailed quantitative characterization of morphological changes in neointima formation, that occurs in vivo, by single-spheroid analysis of various drug treatments.

## INTRODUCTION

Vascular smooth muscle cell (VSMC) proliferation is critical to many biological processes in vascular biology and pathology. VSMC accumulation and proliferation occur during normal vascular development and repair of vascular injury[1-3]. However, pathologies arise when VSMCs fail to cease proliferation once vascular remodeling has been completed. Uncontrolled VSMC accumulation and proliferation, the main pathologic features of vascular injury, restenosis, and atherosclerosis, contribute to intimal thickening or neointima formation[1, 4-12]. Thus, understanding how to control VSMC accumulation and proliferation would advance the effort to treat vascular and cardiovascular diseases. However, clinically effective drug targets for the prevention and treatment of restenosis and neointima formation have not been well established. Moreover, therapeutic responses are frequently different among patients with the same vascular condition. This can be attributed to the heterogeneity of underlying VSMC biology. Chappell et al. reported that only a small subpopulation of the VSMCs at the site of a vascular injury undergo the characteristic phenotypic switch to a synthetic state and actively contribute to hyperplasia resulting in the neointima formation[6]. The lack of a homogenous response poses a significant challenge in formulating patient-specific treatments for vascular diseases, due to current methodologies focusing on characterizing average behaviors of entire cell populations, and thus are limited in detecting such heterogeneity[13-16]. In this paper, we present an experimental and computational pipeline called HETEROID (Heterogeneous Spheroid), which combines a VSMC spheroid model and machine learning (ML) analysis (Fig. 1) to deconvolve the heterogeneous drug effects on VSMC functions involved in neointima formation.

**Figure 1:**
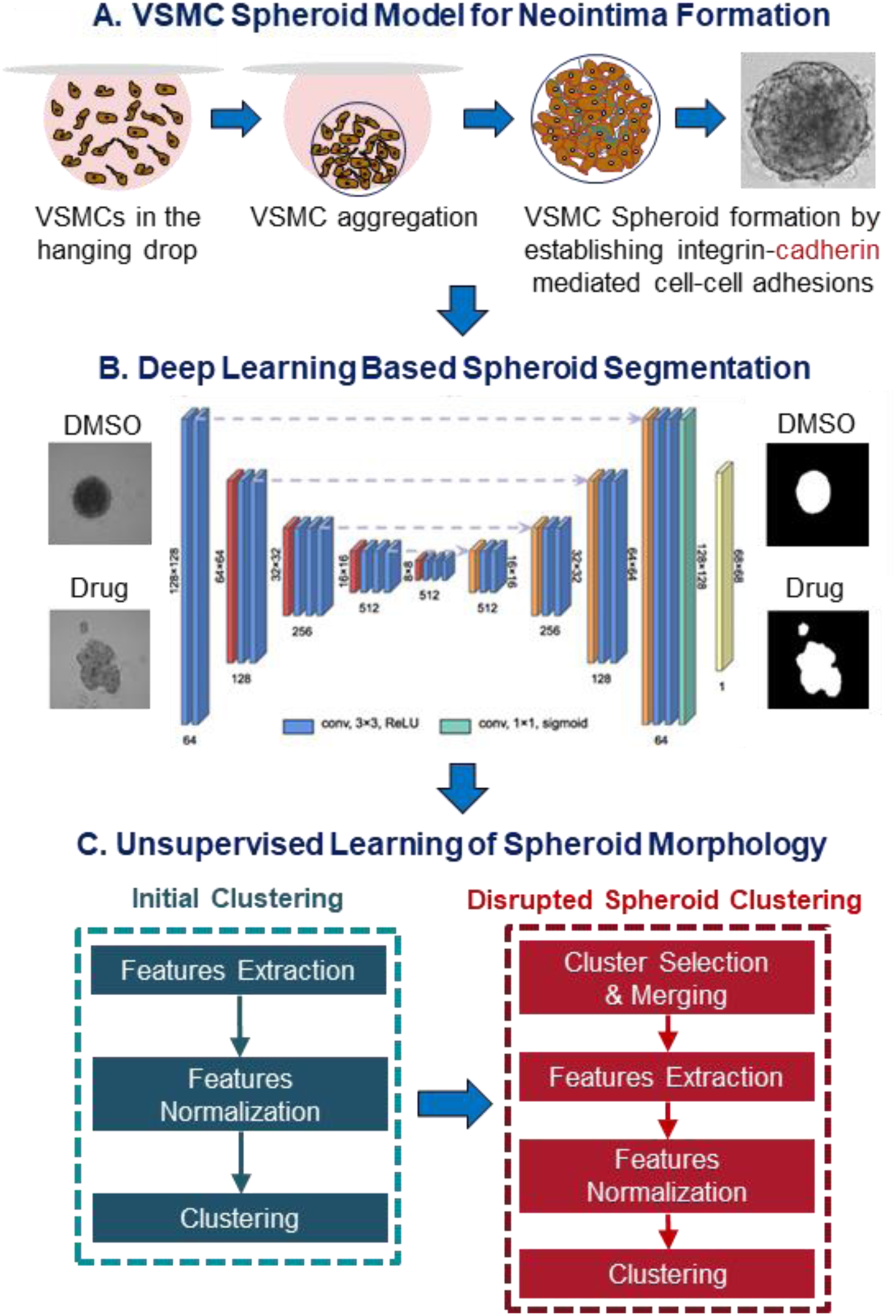
Overview of HETEROID framework: VSMC spheroid images (A) were segmented by VGG19-U-Net semantic segmentation and feature vectors (B) were extracted for additional clustering analysis.(C) Two-level framework was used to analyze the presence of morphological subpopulations of VSMC spheroids. Initial clustering analysis is focused on the roundness of the spheroids. Subsequent clustering analysis is focused on disrupted spheroids.

VSMC interactions among cells (cell-cell adhesion), and between cells and the surrounding extracellular matrix (ECM; cell-ECM adhesion) contribute to neointima formation. How do VSMCs integrate and promote the process of neointima formation? Integrin signaling may facilitate the cell-cell and cell-ECM adhesion as well as cell proliferation during the formation of neointima[17]. The signal transduction by integrin induces the activation of many major focal adhesion proteins including focal adhesion kinase (FAK)[18]. FAK controls cell-cell adhesion and proliferation of VSMCs[11, 12, 19]. Moreover, previous studies reported that FAK is a key component for VSMC-mediated neointima formation following vascular injury[11, 12, 20, 21]. Mui et al showed that FAK activity was increased in neointima after vascular injury in mice[12]. However, in response to vascular injury, VSMC-specific deletion of FAK attenuated the proliferative response and neointima formation through N-cadherin[12]. This study also found that N-cadherin is shown to be upregulated in VSMCs as a result of vascular injury[12]. Further, N-cadherin has been shown to affect cell-cell adhesions as well as cell proliferation[12, 22]. Collectively, FAK and N-cadherin in VSMCs are key signaling molecules that are essential for VSMC-mediated neointima formation.

FAK controls cellular processes including cell adhesion and proliferation by activating its downstream small GTPases Rac, Rho, and Cdc42. First, the activation of Rac by FAK promotes N-cadherin-mediated cell-cell adhesion and cell proliferation[11, 12, 23]. Bond et al., demonstrated that Rac activation is increased following vascular injury in rat, but its inhibition with a dominant-negative Rac1 adenovirus decreased neointima formation[24]. Our previous *in vivo* study also found that VSMC-specific deletion of Rac significantly reduced cell proliferation and neointima formation in response to vascular injury[11, 12]. Moreover, this study showed that Rac inhibition in VSMCs decreased the induction of N-cadherin[12]. Second, Rho activity has been shown to not only promote cell contraction and the formation of stress fiber and focal adhesion but also control VSMC proliferation[25-28]. Furthermore, Rho-associated protein kinase (ROCK) is essential for integrin-mediated cell-cell adhesion and is involved in myosin light chain phosphorylation, which leads to the formation of actin stress fiber and focal adhesions[28]. A previous study showed that Rho inhibition decreased neointima formation and prevented restenosis by increasing VSMC apoptosis after vascular injury[29]. Since FAK regulates Rho activity, FAK can potentially control cell-cell adhesions and neointima formation through Rho. Lastly, FAK also promotes the activation of Cdc42[30-32]. Li et al showed that the inactivation of Cdc42 disrupted N-cadherin-mediated cell-cell adhesion and cell proliferation in cardiomyocytes[32]. Furthermore, endothelial Cdc42 inhibition impaired vascular repair after inflammatory vascular injury[31]. These studies suggest that FAK, Rac, Rho, and Cdc42 are critically involved in the neointima formation. Their differential regulation in VSMC-mediated neointima formation is, however, still not well understood.

In this study, therefore, to test our primary hypothesis that FAK and its downstream targets (Rac, Rho, and Cdc42) are required during the process of spheroid formation, we used VSMC spheroids as a test platform. To do this, we created VSMC spheroids *in vitro*, which are spherical cellular aggregates consisting of multiple layers of VSMC[33, 34], using a hanging drop culture technique[35, 36]. This platform can mimic a VSMC-rich neointima in the vessel wall similar to animal models that simulate human pathologies such as restenosis and atherosclerosis. VSMCs in the spheroid maintain their function and morphology due to cell accumulation through cell-cell adhesions, and internalized signals from the surrounding ECM secreted by the cells[34].

Our VSMC spheroid system was further used to investigate the roles of FAK and its downstream targets in the spheroid formation and morphology following the drug treatment. To extract the morphological parameters from the spheroids, it is necessary to employ an automated image analysis of microscopic images[37-39]. The automated analysis could be used to cluster or group the spheroids based on their morphologies and would serve as a way to study differences in drug responses that may arise amongst spheroids. Several spheroid image analysis platforms have been developed to automatically analyze high-contrast spheroids imaged with brightfield or fluorescence microscopes to quantify spheroid volumes and morphological parameters[40-42]. These softwares are, however, limited in analyzing the morphology of drug-disturbed VSMC spheroids as follows: i) It is very challenging to detect low contrast edges of the disturbed VSMC spheroids from bright-field or phase-contrast images. ii) The existing softwares do not provide the functionality to identify the sub-populations of spheroids that have distinct drug responses. To resolve these issues, we first take a deep learning approach for the spheroid segmentation from phase-contrast images. Specifically, we combine two high-performance deep learning architectures, U-Net and VGG19 pretrained models[43-46]. This transfer learning approach has been widely used to reduce the size of the training set and minimize overfitting[47-50]. After the segmentation, various morphological features were extracted and unsupervised learning was applied to identify distinct subpopulations of VSMC spheroids.

This paper tested the importance of FAK and its downstream small GTPases on VSMC spheroid formation and morphology using our ML pipeline. We found that the inhibition of FAK, Rac, Rho, and Cdc42 disrupted the process of VSMC spheroid formation. Interestingly, inhibition of FAK, Rac, and Rho, but not Cdc42, decreased N-cadherin induction indicating FAK-Rac/Rho pathways control the process of VSMC spheroid formation selectively through N-cadherin. More interestingly, our ML-based clustering analysis computationally identified the morphological variations in drug responses amongst VSMC spheroids. Specifically, we found FAK inhibition disrupted the spheroid morphology and formation more likely through Rac, but not Rho and Cdc42. Moreover, we found that Cdc42 inhibition disrupted the spheroid formation independent of FAK-mediated signaling pathway. Taken together, our ML approach revealed distinct responses to FAK, Rac, Rho, and Cdc42 inhibition on VSMC spheroid formation and morphology. Our ML pipeline can be used to precisely and quantitatively assess the effects of various drugs on the VSMC spheroid model for better characterization of morphological changes in neointima formation that occurs in vascular and cardiovascular diseases.

## RESULTS

### Effect of FAK, Rac, Rho, or Cdc42 inhibition on the process of VSMC spheroid formation

Since FAK is critical in the regulation of cell-cell adhesion, cell-ECM interaction, and cell proliferation during the process of neointima formation[11, 12, 18, 19, 51], we first tested the effect of FAK inhibition on VSMC spheroid formation *in vitro*. To do this, we cultured human VSMCs (Fig. 2A-C) and mouse VSMCs (Fig. S1) to form spheroids in the presence of either PF573228 (a potent and selective FAK inhibitor)[11] or DMSO (a vehicle control). We first confirmed that VSMC spheroid morphologies with DMSO are mostly spherical (Fig. 2C, left panel) as previously shown in other studies[33, 34]. FAK activity (FAK phosphorylation at Tyr 397) in both human VSMCs (Fig. 2A-B) and mouse VSMCs (Fig. S1A-B) was reduced approximately 70-80% with 10 μM PF573228 treatment. The spheroid images acquired at 24 hours showed FAK inhibition significantly disrupted the spheroid formation of both human VSMCs (Fig. 2C, right panel) and mouse VSMCs (Fig. S1C, right panel). Collectively, FAK is required for the process of normal VSMC spheroid formation.

**Figure 2:**
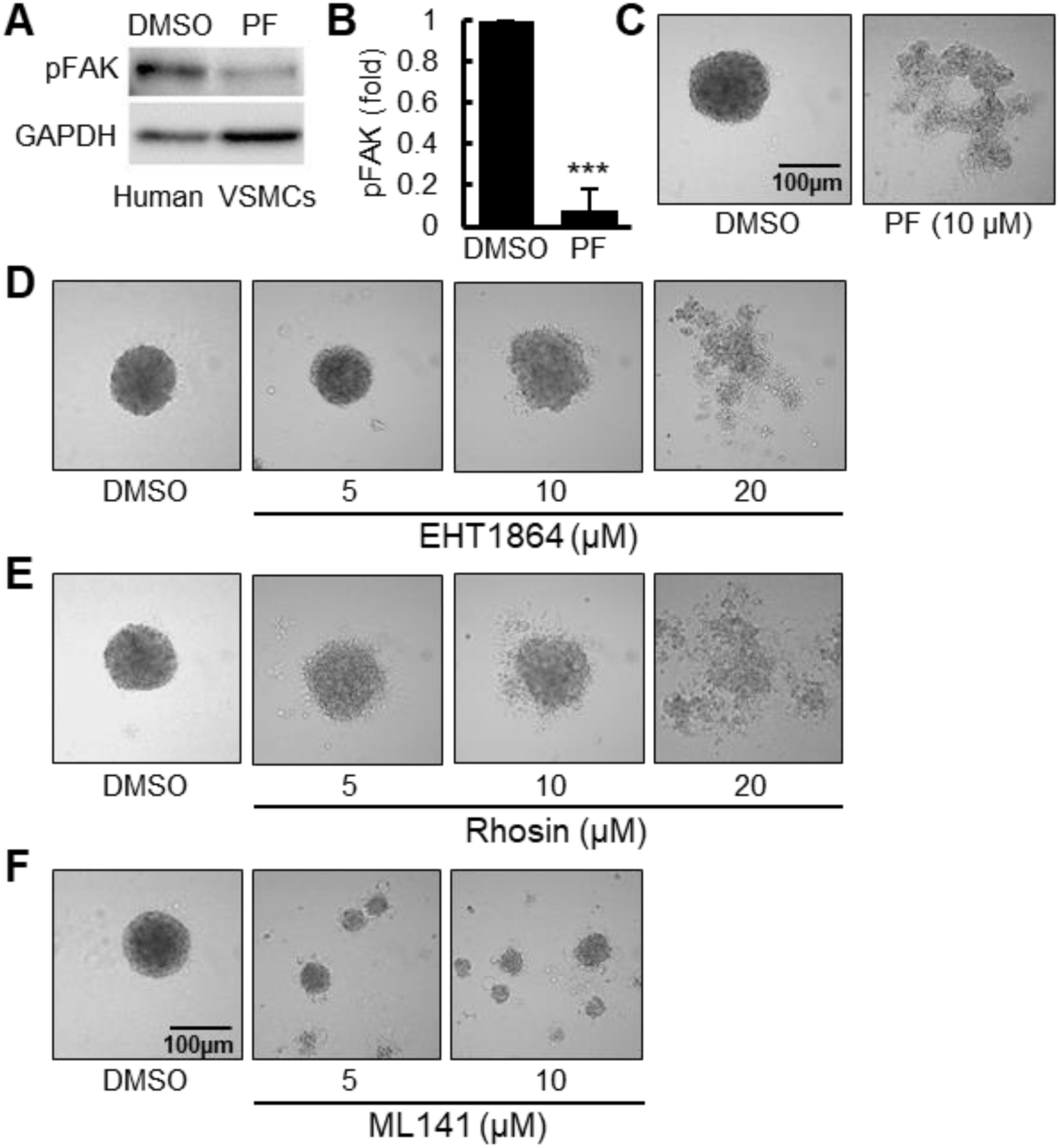
FAK, Rac, Rho, and Cdc42 inhibition disrupt VSMC spheroid formation and morphology. Human VSMC spheroids were made using a hanging drop technique with 2000 cells per droplet in the presence of DMSO (vehicle control), (A-C) PF573228 (PF, FAK inhibitor), (D) EHT1864 (Rac inhibitor), (E) Rhosin (Rho inhibitor), or (F) ML141 (Cdc42 inhibitor) in high glucose DMEM containing 10% serum. Total cell lysates were immunoblotted (A) for phosphorylated FAK at Tyr397 (pFAK) and GAPDH. The bar graph shows pFAK levels normalized to DMSO control (B). Cultures were imaged after 24 hours of incubation using an upright microscope (C and F). *n*=4 (A), *n*=7 (C), *n*=4 (D), *n*=4 (E), and *n*=4 (F) biological replicates were used. *p*\*\*\*<*0*.*001*.

**Supplementary figure 1:**
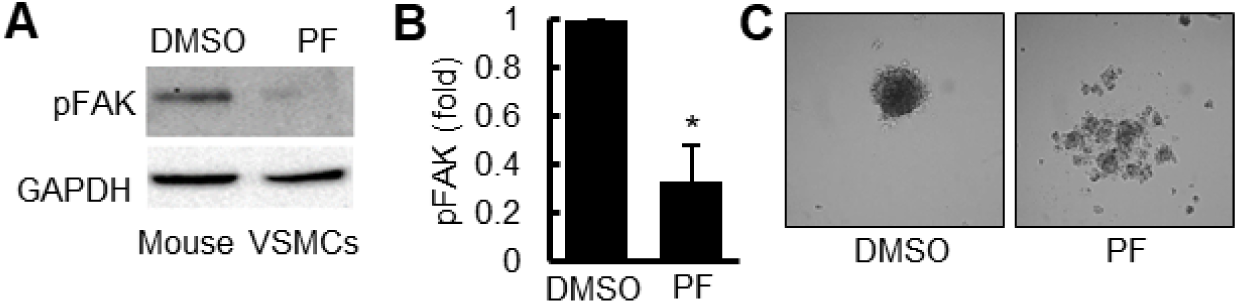
FAK inhibition disrupts mouse VSMC spheroid formation. Mouse VSMC spheroids were made in the presence of either DMSO (vehicle control) or 10 µM PF573228 (PF) in high glucose DMEM containing 10% serum. Total cell lysates were immunoblotted (A) for phosphorylated FAK at Tyr397 (pFAK) and GAPDH. The bar graph displays pFAK levels normalized to DMSO control (B). (C) Cultures were imaged after 24 hours of incubation using an upright microscope. *n*=3 (A) and *n*=12 (C) biological replicates were used. \**p*<*0*.*05*.

Rac, Rho, and Cdc42 are major downstream targets of FAK signaling that regulate cell morphology and cell-cell adhesion[52]. Furthermore, Rac, Rho, and Cdc42 are essential for VSMC proliferation, vascular remodeling, and neointima formation[11, 29, 31, 53]. Therefore, we next evaluated the effects of Rac, Rho, and Cdc42 on spheroid formation by culturing human VSMCs in the presence of 5 to 20 μM EHT1864 (a potent Rac family GTPase inhibitor)[54], 5 to 20 μM Rhosin (a potent, specific inhibitor of RhoA subfamily Rho GTPases)[55], 5 to 10 μM ML141 (an inhibitor of Cdc42 GTPase)[56], or DMSO. Treatments with 5 or 10 μM EHT1864 (Fig. 2D, 2^nd^ and 3^rd^ images) and 5 or 10 μM Rhosin (Fig. 2E, 2^nd^ and 3^rd^ images) had less effect on the formation of VSMC spheroids than DMSO, whereas spheroid formation was significantly disrupted with a higher dose (20 μM) of EHT1864 (Fig. 2D, 4^th^ image) or Rhosin (Fig. 2E, 4^th^ image), or 5-10 μM ML141 (Fig. 2F, 2^nd^ and 3^rd^ images) (compared DMSO-treated VSMC spheroids in Fig. 2D-F, 1^st^ images). This indicates that these small GTPases are required for VSMC spheroid formation. Interestingly, inhibition of Cdc42 resulted in the formation of multiple small cell aggregates, distinct from the disrupted morphologies in cultures treated with inhibitors of FAK, Rac, and Rho. Taken together, these data revealed that the differential effects of FAK, Rac, Rho, or Cdc42 inhibition on the spheroid morphology and formation.

### N-cadherin is a potential downstream effector of FAK, Rac, and Rho-mediated VSMC spheroid formation

Since among other cadherins, N-cadherin is a major cadherin and a downstream target of FAK, Rac, Rho, and Cdc42 in VSMCs[11, 12, 23, 28, 32], disrupted VSMC spheroid formation may be due to the weakening of N-cadherin-mediated cell-cell adhesion, leading to a reduction in VSMC accumulation. In addition, previous studies showed that N-cadherin expression is strongly upregulated in the neointima and underlying tunica media of mouse femoral arteries following vascular injury[12, 22]. However, the injury responses, such as VSMC proliferation and neointimal formation, were significantly decreased in the absence of N-cadherin in VSMCs[12]. Thus, we hypothesized that N-cadherin mediates FAK, Rac, Rho, and/or Cdc42 signals into VSMC spheroid formation. Immunoblot showed a significant reduction in N-cadherin expression with FAK inhibition with PF573228 as compared to control spheroids (Fig. 3A). In spheroids treated with 20 μM EHT1864 (Fig. 3B) and 20 μM Rhosin (Fig. 3C), approximately 50% reduction of N-cadherin expression was observed. However, inhibition of Cdc42 with 5 and 10 μM ML141 did not decrease N-cadherin expression (Fig. 3D). These results indicated that FAK-Rac or FAK-Rho signaling pathways control the process of VSMC spheroid formation possibly through N-cadherin. Taken together, we showed that our VSMC spheroid model could potentially mimic FAK-mediated neointima formation via N-cadherin.

**Figure 3:**
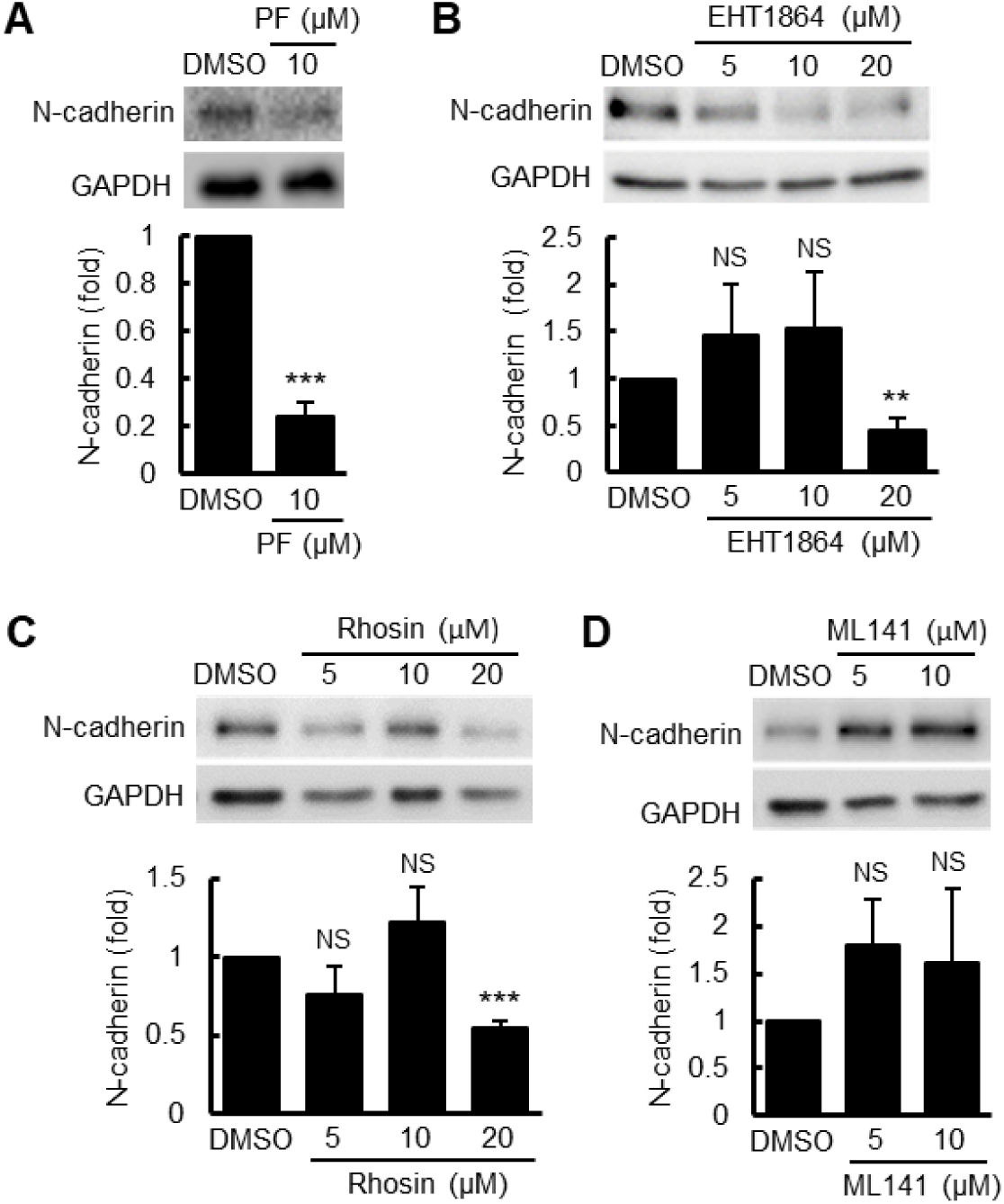
Inhibition of FAK, Rac, and Rho, but not Cdc42, decreased N-cadherin induction. Human VSMCs treated with (A) 10μM PF573228, (B) 5-20 μM EHT1864, (C) 5-20 μM Rhosin, (D) 5-10 μM ML141, or DMSO were used to generate human VSMC spheroids. Total cell lysates were immunoblotted for N-cadherin and GAPDH. *n*=5 biological replicates were used. *p*\*\*<*0*.*05 or p*\*\*\*<*0*.*001*.

### Image segmentation of VSMC spheroid images using deep learning

Interestingly, we observed that FAK, Rac, Rho, and Cdc42-inhibited human VSMC spheroids (Fig. 4) and FAK-inhibited mouse VSMC spheroids (Fig. S2) displayed substantially heterogeneous morphologies from relatively unperturbed to clearly disrupted in both human and mouse VSMC spheroids. These observations suggested the existence of some distinctive characteristics (morphological subpopulations) among VSMC spheroids in response to the same and different drug treatments.

**Figure 4:**
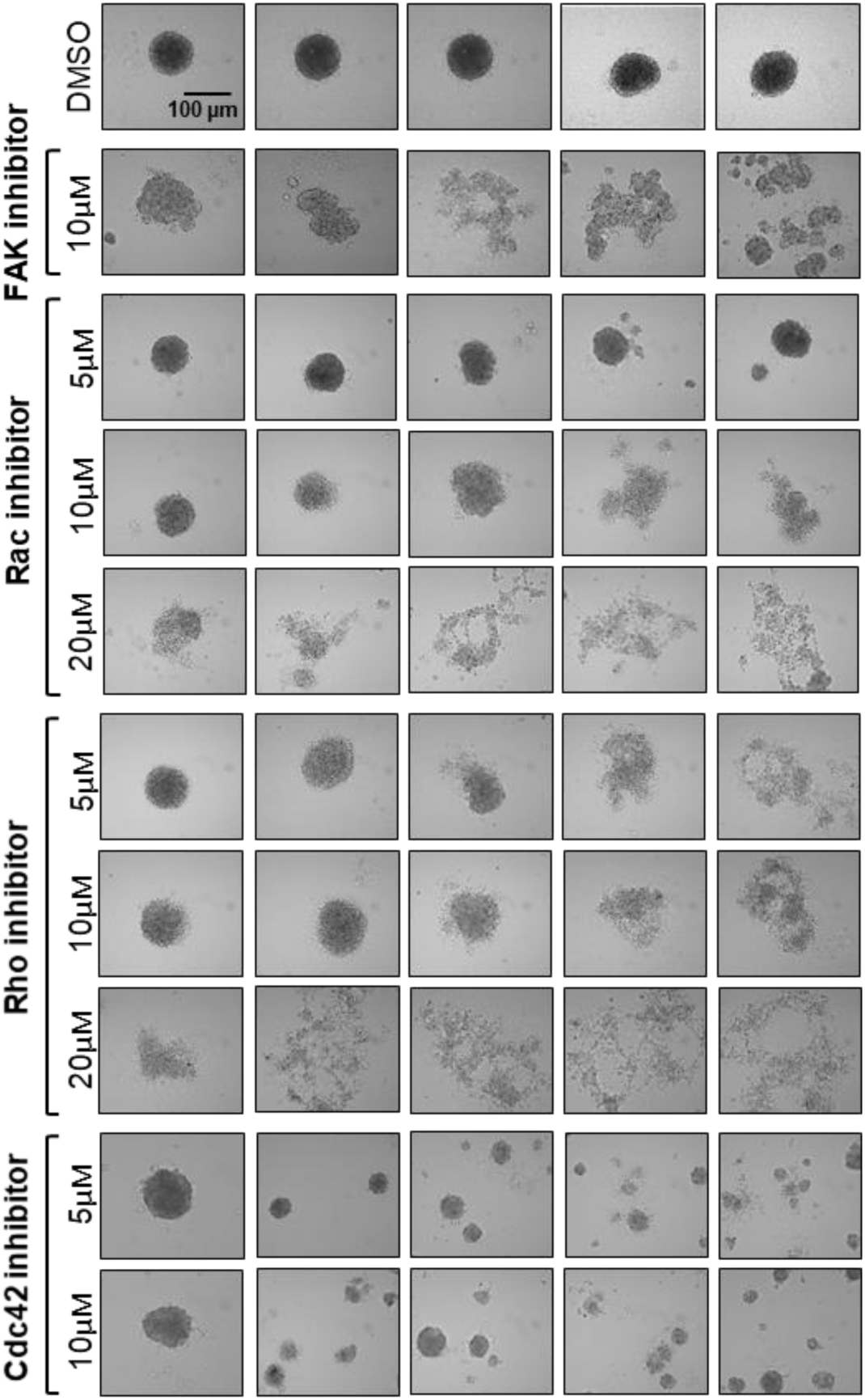
FAK, Rac, Rho, and Cdc42 inhibition differentially affect VSMC spheroid formation and morphology. Human VSMCs treated with inhibitors of FAK (PF573228), Rac (EHT1864), Rho (Rhosin), or Cdc42 (ML141), and DMSO (vehicle control) were used to generate human VSMC spheroids. Spheroids were imaged after 24 hours of incubation using an upright microscope.

**Supplementary figure 2:**
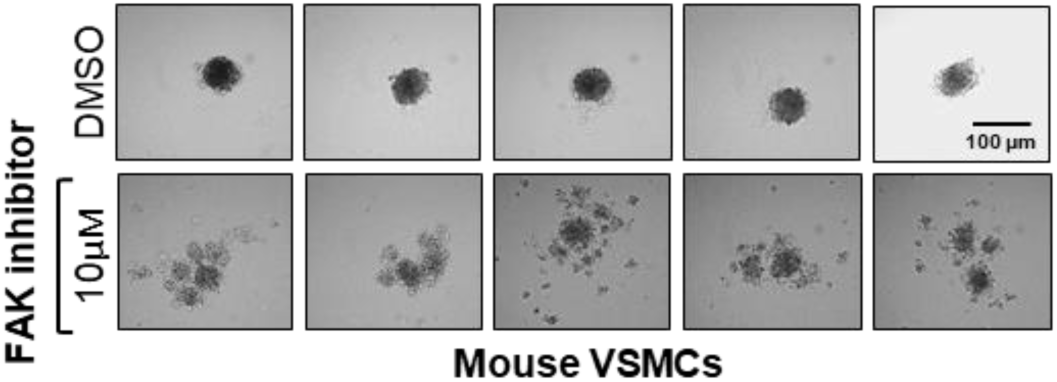
FAK inhibition differentially disrupts mouse VSMC spheroid formation and morphology. Mouse VSMC spheroids were made in the presence of either DMSO (vehicle control) or 10 µM PF573228. Cultures were imaged after 24 hours of incubation using an upright microscope. FAK inhibition shows heterogeneous patterns of disruption in mouse VSMC spheroids. *n*=12.

To speculate on morphological subpopulations of VSMC spheroids following the drug treatment, we further examined changes in spheroid morphology using a machine-learning (ML) based image segmentation, followed by data clustering analysis to identify the presence of morphological subpopulations or clusters resulting from the drug treatments. Since our spheroid images were from a phase-contrast microscope, conventional image segmentation is limited for robust edge detection, particularly for drug-treated spheroids. Therefore, we used a well-known deep learning structure, U-Net integrated with the pretrained VGG19 model, called VGG19-U-Net(Wang, 2019 #56) (Figs. 1B and 5A) to train the deep neural network to segment VSMC phase-contrast images. First, spheroid boundaries were manually drawn on a subset of spheroid images using the Pixel Annotation Tool, and these annotated images were then used to train VGG19-U-Net. When tested with new images outside the training set, the trained VGG19-U-Net identified spheroid boundaries with 95% accuracy at the epoch 30 as demonstrated in the learning curve plotted as dice-coefficient (Fig. 5B) and training loss (Fig. 5C). The VGG19-U-Net produced significantly higher accuracy of the segmentation in comparison to the standard U-Net (Fig. 5D) The images visually confirmed that the algorithm was trained to produce excellent segmentation performance (Fig. 5E-F).

**Figure 5:**
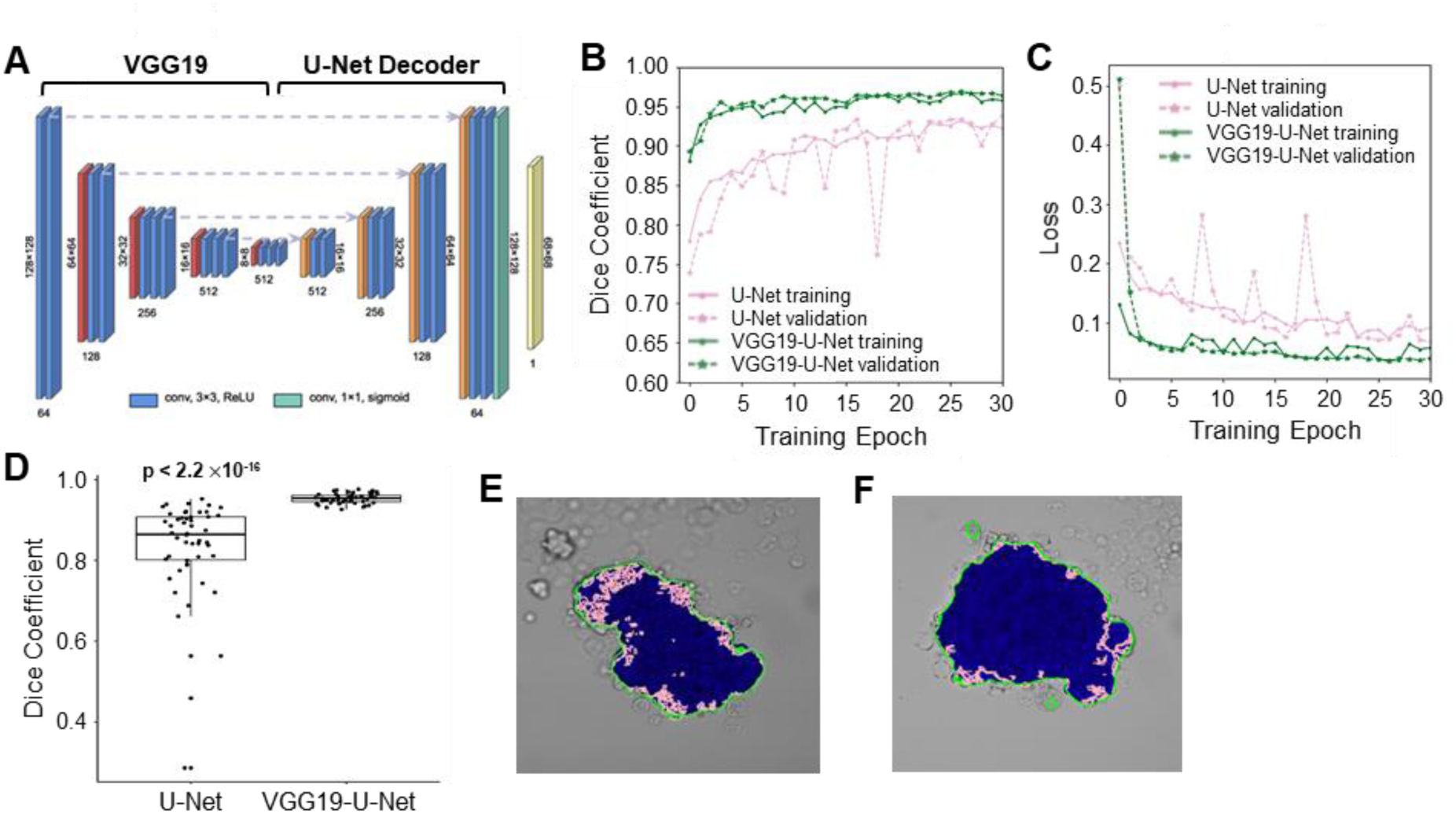
VGG19-U-Net based image segmentation and presence of morphological heterogeneity in response to drug treatments. (A) VGG19-U-Net structure. The training and validation performance of Dice Coefficient (B) and Loss function (C). (D) Comparison of Dice Coefficient between U-Net and VGG19-U-Net. The p-value was calculated by Wilcoxon rank-sum test. (E-F) The edges of the spheroid images identified by U-Net (pink) and VGG19-U-Net (green). The blue regions represent the manually labeled ground-truth.

### Clustering analysis reveals morphological subpopulations

The morphological features of the spheroid images were extracted from the segmented binarized images without considering the grayscale information in the raw images. Five morphological features (eccentricity, solidity, extent, circularity, and the number of colonies; see Methods for detail), were selected and displayed in a UMAP plot, a dimensionality reduction technique[57] (Fig. 6A). The data points scattered on the UMAP plot revealed the presence of morphological variations among spheroids with drug treatments.

**Figure 6:**
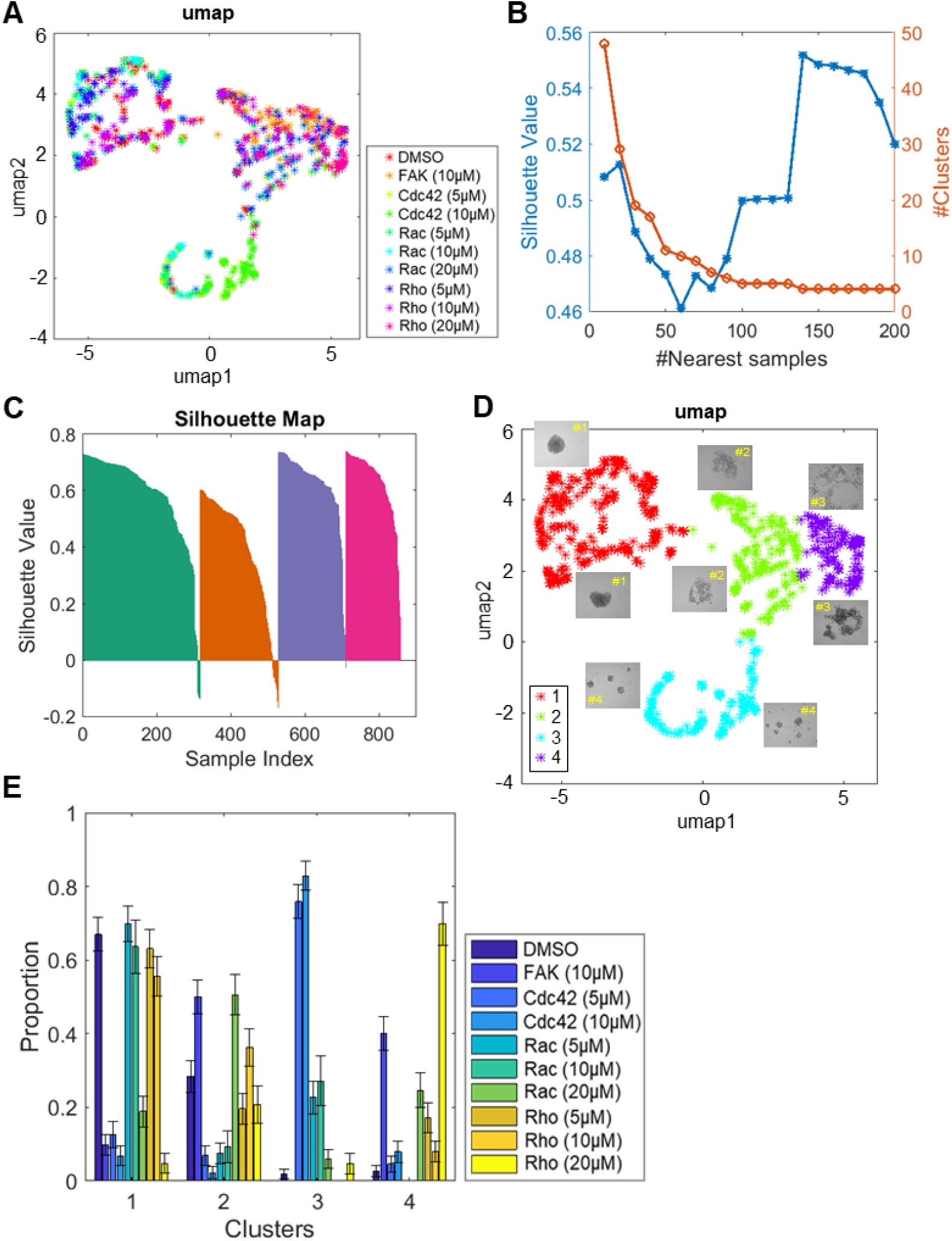
Initial morphological clustering analysis identifies distinct clusters in responses to inhibitors of FAK, Rac, Rho, and Cdc42. (A) Four morphological features extracted from human VSMC spheroid images were plotted as UMAP plot. Different colored ‘*’s are used to represent the various drug treatments; FAK (PF573228), Rac (EHT1864), Rho (Rhosin), and Cdc42 (ML141). (B) The number of nearest samples between 2 and 200 were evaluated and the silhouette values were plotted. A cluster number of four was selected. (C) Silhouette plots for the clustering results. (D) Four morphological clusters plotted on the Umap were observed based on the “roundness of the spheroid”. Different colored ‘*’s are used to represent the various morphological clusters with treatments. The most representative images were inserted into each cluster (2 images per cluster) in the UMAP. Clusters 1 and 3 showed spheroids with circular morphology and the spheroids in clusters 2 and 4 showed disrupted/dispersed or irregular morphologies. (E) A proportionality plot was graphed to identify the distribution of spheroids in each morphological cluster with treatments.

The morphological feature vectors of VSMC spheroids were further analyzed using clustering analysis with the Community Detection algorithm[58] to identify the distinct groups of the spheroid images of similar morphological features. We varied the number of nearest samples, a hyperparameter in the Community Detection algorithm to produce the clustering outcomes of different numbers of clusters. We calculated the quality of each clustering result by evaluating the silhouette value, a measure of how similar the morphology of a given spheroid is to its own cluster compared to other clusters. The highest silhouette value determined the number of clusters and the final clustering result (Fig. 6B). The cluster number of four was chosen based on the maximum silhouette values for the number of nearest samples between 2 and 200. The silhouette plot of the final clustering analysis showed the least overlap between clusters and mostly positive silhouette values (Fig. 6C). The distribution of spheroid morphologies in each cluster is displayed on a UMAP plot (Fig. 6D) and the proportion of spheroids in each cluster for given drug treatment is represented in Fig. 6E. Cluster #1 has morphologies stereotypical of unperturbed circular spheroids (Fig. 6D, inset). Clusters #2 and #4 morphologies were non-circular (dispersed or disrupted). In Cluster #3, images showed small circular shapes but contained more than one spheroid.

### Different drug treatments produced spheroid populations with different morphology profiles

For DMSO-treated controls, most (approximately 70%) were in Cluster (or called Type) #1 (Fig. 6E), as were human VSMC spheroids treated with the inhibitor concentrations (5 or 10 µM Rac and Rho inhibitors, Fig. 6E) that did not reduce N-cadherin expression (Fig. 3). These results indicated that the spheroids in this cluster maintained circular morphology or had minimal effects with DMSO or lower concentration of Rac and Rho inhibitors. In contrast, the proportion of human VSMC spheroids treated with 10 μM FAK inhibitor was reduced to 10% in Cluster #1 and increased up to about 50% in Cluster #2 and 40% in Cluster #4 (Fig. 6E). A high dose (20 μM) treatment of Rac inhibitor produced spheroids that are 50% Type 2 and 25% Type 4. A high dose (20 μM) of Rho inhibition led to producing spheroids that are 20% Type 2 and 70% Type 4, which are significantly fewer and more than FAK and Rac inhibition, respectively. Interestingly, the treatment with 5 or 10 μM Cdc 42 inhibitor created spheroids that are approximately 80% Type 3. Though the spheroids in Type 3 showed circular morphology, each culture contained multiple small spheroids. The morphologies of cultures in which FAK and Rac (20 μM) were inhibited, spheroids were mostly Types 2 and 4 (non-circular and dispersed/disrupted), Rho (20 μM)-inhibited spheroids were mostly Type 4, and Cdc42-inhibited spheroids were mainly Type 3 (small fragmented). Interestingly, however, the data as shown in Fig. 6E also demonstrated the different morphologies of spheroid were generated within a single treatment (i.e. FAK inhibition created approximately 10% Type, 50% Type 2, 2% Type 3, and 40% Type 4 spheroids). Collectively, these results suggest that there exist morphological subpopulations of spheroid formation differentially responding to the drug treatments, suggesting different cellular states (i.e. proliferative and metabolic) following the drug treatment present that affects the process of VSMC spheroid formation.

### Cluster analysis of non-circular spheroids revealed the differential effects of FAK, Rac, and Rho inhibition

The morphological features used in the previous analysis were focused on differentiating circular vs non-circular spheroids, which are limited in distinguishing fine-grained clusters among the non-circular spheroids in Clusters #2 and #4. For a more precise analysis of the spheroids treated with the inhibitors of FAK, Rac (20 μM), and Rho (20 μM) shown in Clusters #2 and #4, we performed further morphological clustering, using a wide range of morphological features (Fig 1C, refers to “Disrupted Spheroid Clustering”). Fifteen features were selected to represent the morphologies of non-circular and disrupted spheroids including area, major axis length, minor axis length, eccentricity, solidity, extent, perimeter, convex area, circularity, aspect ratio, actual area, actual equivalent diameter, actual circularity, and the number of holes. The morphological features extracted from the spheroids from the disrupted non-circular spheroids were plotted on a UMAP (Fig. 7A). Subsequently, clustering analysis with the Community Detection algorithm was performed and the cluster number of four was determined by the maximal silhouette value (Fig. 7B). The silhouette plot of the clustering outcome (Fig. 7C) showed mostly positive and very few negative values indicating the presence of four distinct clusters. The four clusters of drug-treated spheroids with the representative images were plotted on the UMAP plot (Fig. 7D). The spheroids treated with 20 μM Rac inhibitor were almost evenly spread among the disrupted clusters, Types N1-4, while most of human VSMC spheroids treated with 10 μM FAK inhibitor were Types N1, N2, and N4, not in N3, suggesting that Type N3 morphology arises when Rac is inhibited, but not FAK (Fig. 7E). Intriguingly, Type N3 was enriched (50%) when cells were treated with 20 μM Rho inhibitor, while Type N1 was depleted (5%). Overall, we identified VSMC spheroids’ morphological clusters with unique drug responses suggesting the following: i) Types N1 and N4 mostly depend on FAK and Rac. ii) Type N3 mainly depends on Rho. iii) Type N2 depends on FAK, Rac, and Rho (Fig. 7E). iv) Cluster #3 from the previous analysis shown in Fig. 6E depends only on Cdc42.

**Figure 7:**
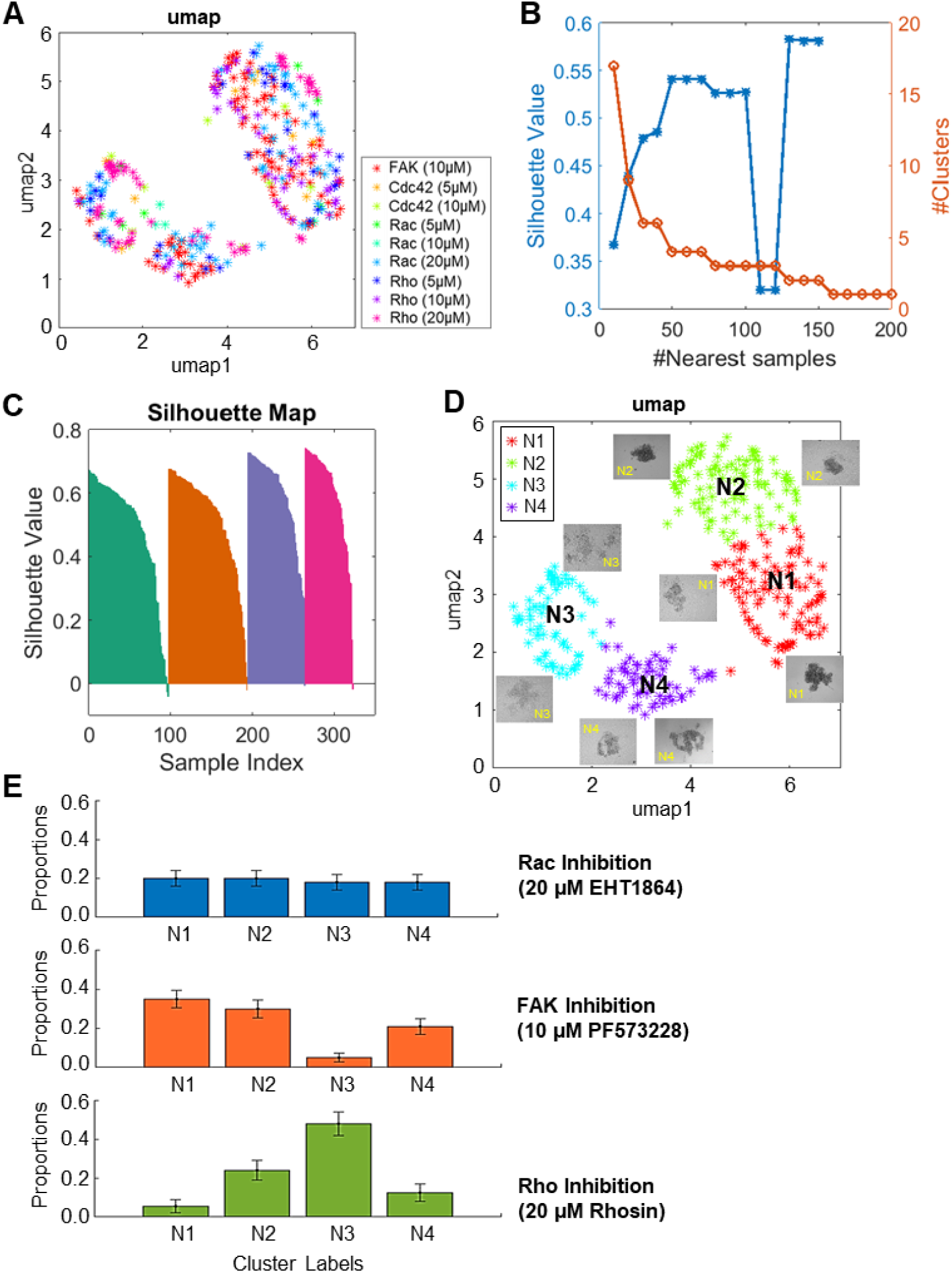
Clustering analysis of non-circular spheroids and the presence of distinct clusters with disrupted morphology. (A) Fifteen morphological features extracted from hVSMC spheroid images from clusters 2 and 4 were graphed as UMAP plot. Different colored ‘*’s are used to represent the various drug treatments. (B) The number of nearest sample between 2 and 200 were evaluated and the silhouette values were plotted. Cluster number of 4 was selected. (C) Silhouette plots for the clustering results. (D) Four morphological clusters were observed based on the non-circular spheroid from Cluster #2 and #4 obtained from Fig. 6. Different colored ‘*’s are used to represent the various morphological clusters. (E) Proportionality plots were plotted to identify the distribution of spheroids in each morphological cluster.

## DISCUSSION

Atherosclerosis and vascular injury are characterized by neointima formation caused by VSMC accumulation and proliferation within the vessel wall[1, 4, 5, 9-12]. In response to vascular injury, VSMCs transition from a highly differentiated state to a de-differentiated state, in which they exhibit increased proliferation, and constitute a substantial part of the neointima[1-3]. While this process resolves smaller insults to the vessel wall, its persistence in chronic injury becomes pathological, generating cellular occlusions in the vessel wall, reducing the blood flow, and creating the need to resolve these changes. Thus, understanding how to control VSMC accumulation and proliferation would advance the effort to treat vascular disease.

Several studies have demonstrated that FAK and its downstream small GTPase proteins (Rac, Rho, and Cdc42) are key regulators of VSMC proliferation, mostly by using a 2D cell culture system and few animal studies. However, the role of FAK and its downstream small GTPase proteins on cell-cell adhesion, cell accumulation, and cell proliferation in a 3D VSMC model has not been explored. In this study, we examined the effects of FAK and downstream small GTPase proteins on VSMC spheroid formation and found heterogeneous cellular responses to the drug treatment. We first showed that FAK inhibition with PF573228 led to marked disruption during the process of VSMC spheroid formation and significantly reduced N-cadherin induction suggesting FAK is required for VSMC accumulation and spheroid formation through N-cadherin-mediated cell-cell adhesion. Moreover, our VSMC spheroid system confirmed the previous findings showing that FAK-N-cadherin signaling is required for forming and maintaining cell-cell adhesions and neointima formation following vascular injury[12, 22].

FAK potentially regulates the process of VSMC spheroid formation through downstream GTPase proteins Rac, Rho, and Cdc42. Thus, we tested the role of Rac, Rho, and Cdc42, known to be major regulators of cadherin-mediated cell-cell adhesions, in VSMC spheroid formation to determine how the process of spheroid formation is disrupted upon FAK inhibition. We observed that inhibition of Rac and Rho with high dose drug treatment, but not Cdc42 inhibition, caused disrupting the process of VSMC spheroid formation with a significant reduction in N-cadherin expression that led to the loss of cell-cell adhesion. Our data can be supported by several previous studies. A study using mesenchymal stem cells showed that N-cadherin mediated cell-cell adhesion and expression is modulated by Rac signaling[59]. Moreover, our previous studies showed that there is an increase in Rac activity after a fine-wire vascular injury to the intimal layer[12]. VSMC-specific deletion of Rac and N-cadherin significantly reduced neointima formation in response to vascular injury[12]. Feil et al. showed that cadherin dependent cell-cell contacts destabilized when endogenous Rho was inhibited[60]. FAK regulates Rho activity through Src and we have shown that FAK can potentially regulate cell-cell adhesions and spheroid formation through Rho. Collectively, our results suggested signaling pathways where FAK controls the process of VSMC spheroid formation through FAK-Rac-N-cadherin or FAK-Rho-N-cadherin pathways.

The direct connection of Cdc42 to VSMC-mediated neointima formation following vascular injury or atherosclerosis remains largely unexplored. Here, we showed that Cdc42 inhibition using ML141 disrupted the formation of the spheroid but generated multiple small spheroids. Unlike Rac and Rho, we found that Cdc42 inhibition resulted in disruption of spheroid formation with no significant changes in N-cadherin levels. This result can be explained by the fact that the inhibition of Cdc42 resulted in creating multiple spheroids into small cellular aggregates (as shown in Figs. 2A and 4), thus levels of total N-cadherin expression and functional N-cadherin-mediated cell-cell adhesion within each small spheroids were rescued and restored, respectively. Moreover, Cdc42 is known to regulate attachment of actin fibers to the sites of cell-cell adhesion that is stabilized by the junctional proteins. Thus, inhibition of Cdc42 could have resulted in a reduction in levels of other adherent junction proteins such as occluding and desmosomes other than N-cadherin[52] which might have resulted in disruption of the spheroids.

A salient result analyzed by our HETEROID pipeline is to computationally identify the presence of heterogeneous morphologies (morphological variations) of the spheroids resulting from the drug treatment with FAK, Rho, Rac, and Cdc42 inhibitors. We used a VGG19-U-net segmentation, which combined the use of U-Net and VGG19 based image segmentation algorithms[44]. Unlike simple ML algorithms that use binary thresholding or filtering, deep learning is designed to learn more in-depth pixel information to identify cell edges (or boundaries). This is particularly useful for phase contrast microscopy which is more suitable to high-throughput applications than fluorescence microscopy. Indeed, our unsupervised ML combined with the clustering analysis approach identified that FAK, Rac, Rho, and Cdc42 inhibitors differentially affect the spheroid morphology. More specifically, we observed that FAK inhibition disrupted VSMC spheroid formation and morphology more likely through Rac, but not Rho and Cdc42. Furthermore, we identified that Cdc42 inhibition disrupted VSMC spheroid formation and morphology independent of FAK-mediated signaling pathway. These results obtained by our HETEROID framework suggested there exist multiple distinct pathways governing the process of VSMC spheroid formation.

Zanoni et al. demonstrated the various morphologies of tumor spheroids exist that correspond to the viability of the tumor cells within the spheroid[61]. Moreover, a recent study on pancreatic cancer spheroids by Laurent et al. reported that the cells in the multicellular tumor spheroids showed varying effects in response to antiproliferative drugs[62]. This variation in results was attributed to the size of the spheroids and the packing of cells in the spheroids. Thus, with our data obtained from VSMC spheroids and our HETROID framework, further exploration of the molecular and functional significance of these distinct morphological clusters (or types) could result in understanding the morphology of the biological relationship. These further investigations will lead us to a) determine the biological and functional roles of FAK, Rac, Rho, and Cdc42 during the process of VSMC spheroid formation, and b) understand why only a subpopulation of VSMCs underlying the site of vascular injury is actively involved in hyperplasia during the process of VSMC-rich neointima formation[6]. Therefore, these previous studies and our current and future studies will potentially be a development in helping to create specific drug targets for spheroids based on heterogeneous responses in spheroid morphologies.

## CONCLUSION

This study tested the effect of FAK and its downstream small GTPase manipulation on the process of VSMC spheroid formation and used the machine learning approach to reveal heterogeneous responses to FAK, Rac, Rho, and Cdc42 inhibition on VSMC spheroid morphology. The machine learning framework was designed to extract morphological features of the spheroids from bright-field images without using fluorescent staining. Results shown in our study suggest that there is significant drug-induced heterogeneity in spheroid formation and morphology, which has been overlooked in previous analyses and studies. These detected subpopulations upon further study could give rise to the biological significance of various drug responses. For instance, examining the effects of the drug treatment on global changes in genome or proteome and on viability, proliferation, or apoptosis in VSMC spheroids within each morphological cluster (shown in Figs. 6D and 7D) can further reveal biological and functional significance that is associated with changes in spheroid morphology. The biological and functional significance of FAK, Rac, Rho, and Cdc42-mediated spheroid formation and morphologies will be a stepping-stone to identifying potential pharmaceutical drug targets on preventing neointima formation. Overall, this is our first step towards developing an automated method for quickly and precisely analyzing the heterogeneity of VSMC spheroids using machine learning. This machine learning approach can be used to study the effects of different pharmacological drugs on VSMC spheroid models for better characterization of pathophysiologic progression of diseases like atherosclerosis, hypertension, restenosis, etc. The long term goal of the future study is to be able to use machine learning algorithms to assist in personalized medicine by testing the effects of pharmacological drugs on patient-specific spheroids before prescribing the drugs to patients.

## ACKNOWLEDGMENTS

This work was funded in part by American Heart Association-Career Development Award-18CDA34080415 (to Y.B.) and NIH grant R35GM133725 (to K.L.). The funders had no role in study design, data collection and analysis, decision to publish, or preparation of the manuscript.

## AUTHOR CONTRIBUTIONS

K.V., C.W., S.J.H., J.K., K.L., and Y.B. designed research. K.V. and A.K. performed experiments. K.V., C.W., Y.Y., M.C. B.L. performed analysis. K.V., C.W., S.J.H., J.K., K.L., and Y.B. wrote the manuscript. All authors critically reviewed the manuscript.

## DECLARATION OF INTERESTS

The authors declare no competing interests.

## STAR METHODS

### Cell culture and drug treatment

Primary human vascular smooth muscle cells (ATCC) and mouse VSMCs (Cell Biologics) were cultured as previously described[8, 11, 63]. Human and mouse VSMCs were used at passages 3-8 and 3-5, respectively. For serum starvation to synchronize cells, near confluent VSMCs were incubated for 48 hours in serum-free DMEM containing 1 mg/ml heat-inactivated, fatty-acid free bovine serum albumin (BSA; Tocris). For FAK, Rac, Rho, and Cdc42 pharmacologic inhibitor experiments, cells in suspension culture were treated with 10 μM FAK specific inhibitor PF573228 (Sigma)[11], 5-20 μM Rac specific inhibitor EHT1864 (Tocris) [54], 5-20 μM Rho specific inhibitor Rhosin (Tocris)[55], or 5-10 μM Cdc42 specific inhibitor ML 141 (Tocris)[56] reconstituted in Dimethyl sulfoxide (DMSO) for selected times up to 24 hours.

### Generation of VSMC spheroids

VSMC spheroids were generated by the hanging drop approach[35, 36]. Briefly, serum-starved cells were trypsinized, centrifuged, and resuspended in fresh high-glucose (4.5g/L) DMEM containing 10% fetal bovine serum (FBS). Total cell numbers were counted with a hemocytometer. Cell suspension was diluted to have approximately 2000 cells per 20 μL high-glucose DMEM with 10% FBS and treated with either DMSO (vehicle control) or pharmacological inhibitors (PF573228, EHT1864, Rhosin or ML141) before plating. Approximately 20 to 30 cell-suspension droplets were dispensed on the hydrophobic surface of the lid of a 100-mm tissue culture dish. Note that the drops were seeded sufficiently apart to avoid the mixture of each adjacent drop. To prevent evaporation of the drops, a water reservoir (hydration chamber) was made by adding 8 ml of sterile water + 2 ml of DMEM at the bottom of the tissue culture dish. The lid was then carefully inverted so cells remained suspended and were allowed cells to aggregate by gravity. The cultures were maintained at 37°C and 10% (human VSMCs) or 5% (mouse VSMCs) CO_2_. VSMC spheroids formed after 24-48 hours of incubation. Images were taken at 10X magnification using an upright microscope (Olympus BH-2) equipped with a digital camera (AmScope) to monitor spheroid formation.

### Protein extraction and immunoblotting

Total cell lysates were prepared from human and mouse VSMC spheroids. Spheroids were collected in 1.7 ml Eppendorf tubes, centrifuged, and extracted in 1X TNE lysis buffer [50 mM Tris-HCl (pH 8.0), 250 mM NaCl, 2 mM EDTA, 1% Nonidet P-40, plus protease and phosphatase inhibitor cocktail composed of a proprietary mix of AEBSF, aprotinin, bestatin, E64, leupeptin, and pepstatin A to promote broad-spectrum protection against endogenous proteases]. Equal amounts of extracted protein were mixed with 5x sample buffer [250 mM Tris-HCl (pH 6.8), 10% SDS, 50% glycerol, 0.02% bromophenol blue, 10 mM 2-mercaptoethanol], denatured by boiling samples at 100 °C, subjected to 8% sodium dodecyl sulfate (SDS) polyacrylamide gel electrophoresis and transferred to a polyvinylidene difluoride (PVDF) membrane. The PVDF membranes were blocked with either 5% BSA or 5% milk in 1X TBST (Tris-buffered saline, 0.1% Tween 20) for 1 hour and probed with primary antibodies to phospho-FAK (Invitrogen, 44-624G), N-cadherin (Proteintech, 22018-I-AP), and GAPDH (Proteintech, 60004-I-Ig). After overnight incubation at 4°C, the PVDF membranes were washed with 1X TBST and probed with horseradish peroxidase-conjugated secondary antibodies (Bio-Rad) for 1 to 2 hours at room temperature. Immunoblot signals were detected using chemiluminescence (Bio-Rad).

### Deep learning-based segmentation of spheroid images

The detailed training procedure is described elsewhere[16].

#### i) Data labeling

A deep neural network (DNN) was trained to automatically segment spheroid images. For the training of the DNN, a set of twenty VSMC spheroids from each of DMSO and drug-treated groups were randomly selected and spheroid boundaries were manually drawn using the Pixel Annotation Tool software (https://github.com/abreheret/PixelAnnotationonTool). The boundary of each spheroid was traced and the resultant images were used as 61 raw images for the next step. Images of the hand-drawn boundaries were used to generate binary images. The masks were used as a training data set from which the DNN learns to identify the boundary of the spheroids.

#### ii) Pre-processing

For each image, all the pixel values were collected from the labeled frames. Then, we calculated the mean µ and the standard deviation δ of pixel values. We replaced the pixel values *x*_*i,j*_ with the follows values when they are less than µ − 2δ or greater than µ + 3δ.

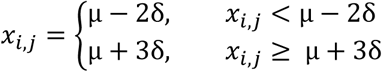

We applied the min-max normalization to rescale the pixel ranges to [0, 255].

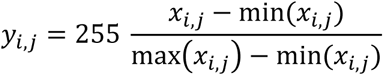

#### iii) Training Dataset Preparation

We randomly cropped 200 patches (128×128 pixels) from each frame. The 60% of the cropped patches contain the edge boundary and 40% contain only to foreground or background. After the image patches were cropped from the whole 20 images, the mean and standard deviation of collected image patches are calculated for further normalization. After preprocessing, we split them into training and validation set with a ratio of 0.8:0.2. Then, the standard data augmentation process was applied using the *ImageDataGenerator* implemented in Keras package. The parameters in the *ImageDataGenerator* for the data augmentation is as follows: *rotation_range*=90; *horitontal_flip*=true; *vertical_flip*=true.

#### iv) Training the deep neural network

The DNN is divided into two parts: an encoder that extracts image features and a decoder that identifies the edge location using the extracted features. The VGG19 model was used in the encoder and extracted features at multiple levels. The initial weights for the VGG19 model were from the pretrained model using ImageNet DB. The VGG19 weights were changed during the training. The decoder from U-Net is incorporated to combine the VGG19 features to generate the segmented images. The generated masks were then overlapped with raw images to visualize the learning efficiency of the algorithm. The binary cross-entropy was used as a loss function for training. Adam was used as an optimizer, and the initial learning rate was 5×10^−4^, and other parameters were default values in the Keras. To avoid overfitting, we used early stopping. We stopped the training when the validation loss did not decrease 0.0001 in consecutive three epochs. The maximum epoch was 30, and the batch size was 32. The neural network training was performed using Keras with TensorFlow backend on NVIDIA GTX 1080Ti or Titan X.

#### v) Performance evaluation of edge localization

We used the labeled data which are not included in the training process (training/validation sets) for the performance evaluation. We generated the binarized mask images by thresholding the softmax output images using the *im2bw* (MATLAB) function with the threshold value of 0.5. Then, we generated edge images using the *bwboundaries* (MATLAB) function. The edge images were predicted for all the image frames including the labeled and unlabeled images. We used the predictions of labeled images not used for training to evaluate the performance of edge localization.

We calculated Dice coefficients to evaluate the segmentation performance along the edge boundary as follows. We applied binary AND operation between the dilated edge masks and the masks of the ground truth and the predicted masks from VGG19-U-Net using the *bitwise_and* function in OpenCV package. After that, we calculate the Dice coefficient between the ground truth and the predictions for each image frame using the following formula (|*A* ∩ *B*| : the common elements between sets *A* and *B*, |*A*| and |*B*| : the number of elements in set *A* and *B*).

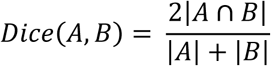

### Unsupervised Learning of Spheroid Morphology

#### i) Feature extraction

The segmented spheroid images were processed for extracting the morphological features. The morphological features were used for unsupervised learning to characterize the responses of spheroids to drug treatments. The morphological features extracted are listed in the following clustering analysis section. Once the features were quantified, feature standardization was used to make each feature in each experiment zero-mean and unit-variance to remove batch effects, followed by Principal Component Analysis. The feature vectors extracted from the spheroids were visualized by UMAP.

#### ii) Clustering analysis

The feature vectors generated from the spheroid images were analyzed to identify the presence of morphological subpopulations. The community detection clustering analysis was performed using the feature vectors in two steps: Clustering of whole spheroids (step 1) and disrupted spheroids (step 2). The number of clusters was determined by the maximum of silhouette values of the clustering results.

a. ***Clustering of whole spheroids (step 1):*** Five morphological features relating to the roundness of the spheroids were used for clustering the spheroids. The features were *Circularity* (4π Area/Perimeter^2^), *Extent* (the spheroid colony with the largest area), *Solidity* (proportion of the pixels in the convex polygon that is also in the region), and *Eccentricity* (an ellipse whose eccentricity is 0 is a circle, while an ellipse whose eccentricity is with *the Number of Colonies*. The resulting clusters were expected to show clusters of spheroid images with round morphology and clusters with disrupted morphologies.
b. ***Clustering of disrupted spheroids (step 2):*** Clusters containing the non-circular spheroids (Clusters #2 and #4; Fig. 6E) were selected and more features were extracted from the selected spheroids for further clustering analysis. Additional morphological features were extracted from the non-circular spheroid clusters. These morphological features were more focused on exploring morphology with various complex patterns of the spheroid. The morphological features extracted include *Area* (area of all the pixels in the largest external contour), *Major axis length* (length in pixels of the major axis of the ellipse that has same normalized second central moments as the region), *Minor axis length* (length in pixels of the minor axis of the ellipse that has same normalized second central moments as the region), *Eccentricity, Solidity, Extent* (ratio of pixels in the region to pixels in the total bounding box, returned as a scalar), *Perimeter* (distance around the boundary of the region returned as a scalar), *Convex area* (number of pixels in the bounding box of the region), *Circularity, Aspect ratio, Actual area, Actual equivalent diameter* (diameter of a sphere with the same volume as the region), *Actual circularity*, and *Number of holes* (number of internal contours).
c. We quantified the drug effect based on the cluster proportion calculating as follows. We counted the number of spheroid images of each cluster for the control and drug treatment experiments. Then the number was resampled using *bootstrp*() in MATLAB to build 10,000 different bootstrapped dataset and the distribution of the proportions in each experiment was generated. Based on these distributions, p-values were calculated by estimating the probability that the cluster proportion was greater or less than that of the other experiment. In addition, 95% confidence interval of the proportions were estimated by *bootci*() functions in MATLAB.

### Statistical analysis

Data are presented as mean + standard error of mean deviation (SE) of the indicated number of independent experiments. Data were analyzed using the student t-test. Results with p-values lower than 0.05 (*), 0.01 (**), or 0.001 (***) were considered to be statistically significant.

## Notes

### Competing Interest Statement

The authors have declared no competing interest.

